# The opposing Chloride Cotransporters KCC and NKCC control locomotor activity in constant light and during long days

**DOI:** 10.1101/2021.09.14.460201

**Authors:** Anna Katharina Eick, Maite Ogueta, Edgar Buhl, James J. L. Hodge, Ralf Stanewsky

**Author notes:** Corresponding author and Lead Contact, Ralf Stanewsky.

## Abstract

Cation Chloride Cotransporters (CCC’s) regulate intracellular chloride ion concentration ([Cl^−^]_i_) within neurons, which can reverse the direction of the neuronal response to the neurotransmitter GABA. Na^+^ K^+^ Cl^−^ (NKCC) and K^+^ Cl^−^ (KCC) cotransporters transport Cl^−^ into or out of the cell, respectively. When NKCC activity dominates, the resulting high [Cl^−^]_i_ can lead to an excitatory and depolarizing response of the neuron upon GABA_A_ receptor opening, while KCC dominance has the opposite effect. This inhibitory-to-excitatory GABA switch has been linked to seasonal adaption of circadian clock function to changing day length, and its dysregulation is associated with neurodevelopmental disorders such as epilepsy. Constant light normally disrupts circadian clock function and leads to arrhythmic behavior. Here, we demonstrate a function for KCC in regulating *Drosophila* locomotor activity and GABA responses in circadian clock neurons because alteration of KCC expression in circadian clock neurons elicits rhythmic behavior in constant light. We observed the same effects after downregulation of the Wnk and Fray kinases, which modulate CCC activity in a [Cl^−^]_i_-dependent manner. Patch-clamp recordings from clock neurons show that downregulation of KCC results in a more positive GABA reversal potential, while KCC overexpression has the opposite effect. Finally, KCC downregulation represses morning behavioral activity during long photoperiods, while downregulation of NKCC promotes morning activity. In summary, our results support a model in which the regulation of [Cl^−^]_i_ by a KCC/NKCC/Wnk/Fray feedback loop determines the response of clock neurons to GABA, which is important for adjusting behavioral activity to constant light and long-day conditions.

## Results

### Altered *kcc* expression in clock neurons leads to abnormal rhythmicity in constant light

We previously reported that the Na^+^ K^+^ Cl^−^ cotransporter encoded by CG31547 and the interacting GPI-anchored membrane protein Quasimodo (Qsm) contribute to normal fly behavior in constant light (LL) ^1,2^. Downregulation or overexpression of either gene led to abnormal rhythmicity in LL, indicating their requirement for normal light-input to the circadian clock. Moreover, simultaneous downregulation of both genes restored wild type behavior, suggesting a genetic interaction. Indeed, downregulation of *qsm* resulted in an elevated firing rate of large ventrolateral clock neurons (l-LNv), which could be restored to wild type levels by parallel knock-down of *NKCC* ^1^. Knock-down of both genes also affected the GABA reversal potential (E_GABA_) in the opposite way, with *qsm-RNAi* resulting in a more positive E_GABA_, leading to occasional excitatory GABA function ^1^. These results indicated that [Cl^−^]_i_ plays an important role in light-dependent behavior, and we therefore set out to test if the K^+^ Cl^−^ cotransporter encoded by *kazachoc* (*kcc*), which in contrast to NKCC transports Cl^−^ out of the cell, also plays a role in light-input to the circadian clock.

For this, we crossed three independent *kcc-RNAi* lines to the *tim-Gal4* driver, which is expressed in all clock cells, including central brain clock neurons and glia, as well as peripheral clock cells ^3^. Locomotor activity of male progeny was tested in LL (~100 lux white light) using the Trikinetics monitoring system ^1^ (STAR Methods). Individual flies were classified either as ‘arrhythmic’ (typical behavior for wild type flies in LL), ‘split’ (> 1 rhythmic component), or circadian (one rhythmic component) as described in Methods (Figure 1A, C). Compared to the control flies (Gal4 and UAS flies crossed to Canton S or *y w*), flies expressing *kcc-RNAi* in all clock cells showed a ~ 40% increase in LL-rhythmicity (Figure 1B, C, Table S1). The relatively high percentage of rhythmic control flies (between 37% and 57%, Table 1) is attributable to the low light intensity (~100 lux white light) used. This is necessary because all flies carry a wild type copy of the *cryptochrome* (*cry*) gene, which is the main photoreceptor responsible for arrhythmic behavior in LL ^4,5^. Nevertheless, the drastic increase in LL-rhythmicity associated with *kcc-RNAi* expression in all clock cells indicates a defective light-input to the circadian clock, and a role for *kcc* in this process. To distinguish between a role for *kcc* in central brain clock neurons and peripheral clock cells we next used the *Clk856-Gal4* driver, which is specifically expressed in all 150 brain clock neurons, but not in glia or peripheral clock cells ^6^. *Clk856-Gal4* crossed to two *kcc-RNAi* lines resulted in a > 30% increase of LL-rhythmicity compared to the controls, while with the 3^rd^ line the effect was less pronounced (> 10% increase, see Figure 1B, C, Table S1). These results do not rule out a contribution of *kcc* function in peripheral clock and glia cells, but they do show that *kcc* expression in central clock neurons is important for circadian light-input.

**Figure 1:**
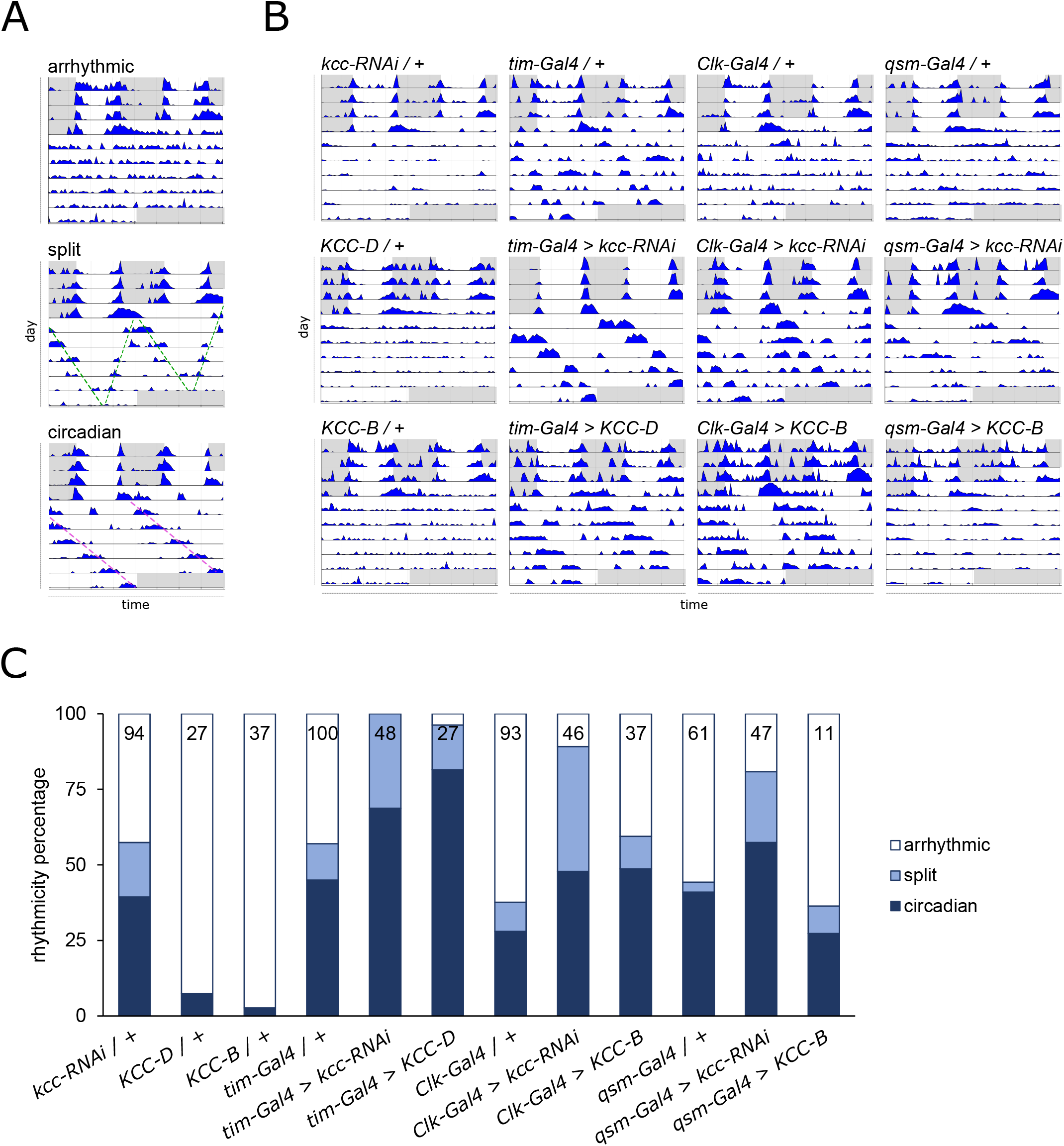
Altered *kcc* expression in clock neurons leads to abnormal rhythmicity in constant light. **(A)** Exemplary double-plotted actograms of individual flies exhibiting arrhythmic, split or circadian behavior under constant light (LL). Locomotor activity was recorded for three days in 12 h : 12 h LD followed by seven days in LL at ~100 lux (gray: lights off; white: lights on) at 25°C. Based on actograms and rhythmicity analysis, flies were manually scored as rhythmic or arrhythmic (see Methods). Rhythmic behavior was further classified as split (complex locomotor rhythm with long and short period; green lines) or circadian (single locomotor rhythm; magenta lines). Genotypes are *Clk856-Gal4 / +* (arrhythmic), *Clk856-Gal4 > wnk-RNAi KK106928* (split) and *tim-Gal4:27 > kcc-RNAi KK101742* (circadian). **(B)** Representative actograms of individual flies with altered *kcc* expression and of individual control flies. Flies express *kcc-RNAi (KK101742), KCC-D* or *KCC-B* under the control of *tim-Gal4:27, Clk856-Gal4* or *qsm^105^-Gal4*. For behavioral controls, *Gal4* drivers and *UAS* lines were crossed to Canton S or *y w*. **(C)** Percentages of flies exhibiting circadian (dark blue), split (light blue) or arrhythmic (white) behavior in LL. Number of flies per genotype given inside bars.

Next, we overexpressed the two major KCC isoforms—encoded by *KCC-B* and *KCC-D*--which differ in their first coding exon and are expressed in the adult brain and during development, respectively ^7^. Expression of the brain-specific isoform *KCC-B* using the *tim-Gal4* driver resulted in flies with drastically reduced life span (flies died after 2-4 days), precluding behavioral analysis. In contrast, overexpression of *KCC-D* with the same driver resulted in flies with normal life span that where strongly rhythmic in LL (96%, Figure 1B, C, Table S1). Overexpression of either isoform in all clock neurons (*Clk856-gal4*) resulted in flies with normal life span, which also exhibited robust rhythmicity in LL (Figure 1B, C, Table S1). These results are similar to a previous report, where both downregulation and overexpression of NKCC resulted in LL-rhythmicity and suggests that disruption of the endogenous [Cl^−^]_i_ balance interferes with normal light-input to the clock (^1^, and see Discussion).

As mentioned above, simultaneous downregulation of NKCC and Qsm restored wild type (i.e., arrhythmic) behavior in LL ^1^. If NKCC and KCC indeed affect Qsm- and light-dependent LL behavior in opposite ways, simultaneous downregulation of Qsm and KCC should result in rhythmic LL behavior. Downregulation of Qsm alone, using one copy of the hypomorphic *Gal4* insertion *qsm^105^* (*qsm^105^/+*) results in a moderate percentage of LL-rhythmicity (44%, Figure 1B, C, Table S1) ^1,2^. We then used this intronic *Gal4* insertion to express *kcc-RNAi* in *qsm* expressing cells ^2^ and observed an increase of LL-rhythmicity with all three RNAi lines, both compared to *qsm^105^/+* (12-37% increase depending on RNAi line), and compared to the different RNAi lines/+ (19% - 42% increase depending on RNAi line) (Figure 1B, C, Table S1). In contrast, overexpression of KCC in the face of *qsm^105^* /+ did not alter the residual LL-rhythmicity of heterozygous *qsm^105^* flies (Figure 1B, C, Table S1). These results therefore support the opposing roles of NKCC and KCC in regulating Qsm-dependent LL behavior.

### The kinases Wnk and Fray are required for normal light-input to the circadian clock

Wnk kinase is inhibited by high [Cl^−^]_i_ and activated by low [Cl^−^]_i_ and active Wnk phosphorylates the Fray kinase. Activated Fray in turn inhibits KCC and activates NKCC by phosphorylation and this feedback loop thus determines [Cl^−^]_i_ ^8^. To test if this feedback loop and [Cl^−^]_i_ are indeed important for LL behavior, we downregulated *Wnk* and *fray* using the *tim-Gal4 and Clk856-Gal4* driver lines. This manipulation is expected to block NKCC and activate KCC function lowering [Cl^−^]_i_ and should therefore result in LL rhythmicity, similar to *NKCC-RNAi* and *kcc* overexpressing flies (Figure 1, Table S1) ^1^. We observed a drastic increase in LL rhythmicity in two independent *wnk-RNAi* lines crossed to both *tim-Gal4* and *Clk856-Gal4* driver lines (Figure 2, Table S1). Similarly, robust LL-rhythmicity was obtained with four *fray-RNAi* lines (Figure 2, Table S1), confirming that both kinases are important for normal light signaling to the clock. These results point to a crucial role of [Cl^−^]_i_, regulated by the NKCC/KCC and Wnk/Fray feedback loop, required for normal LL behavior in *Drosophila*.

**Figure 2:**
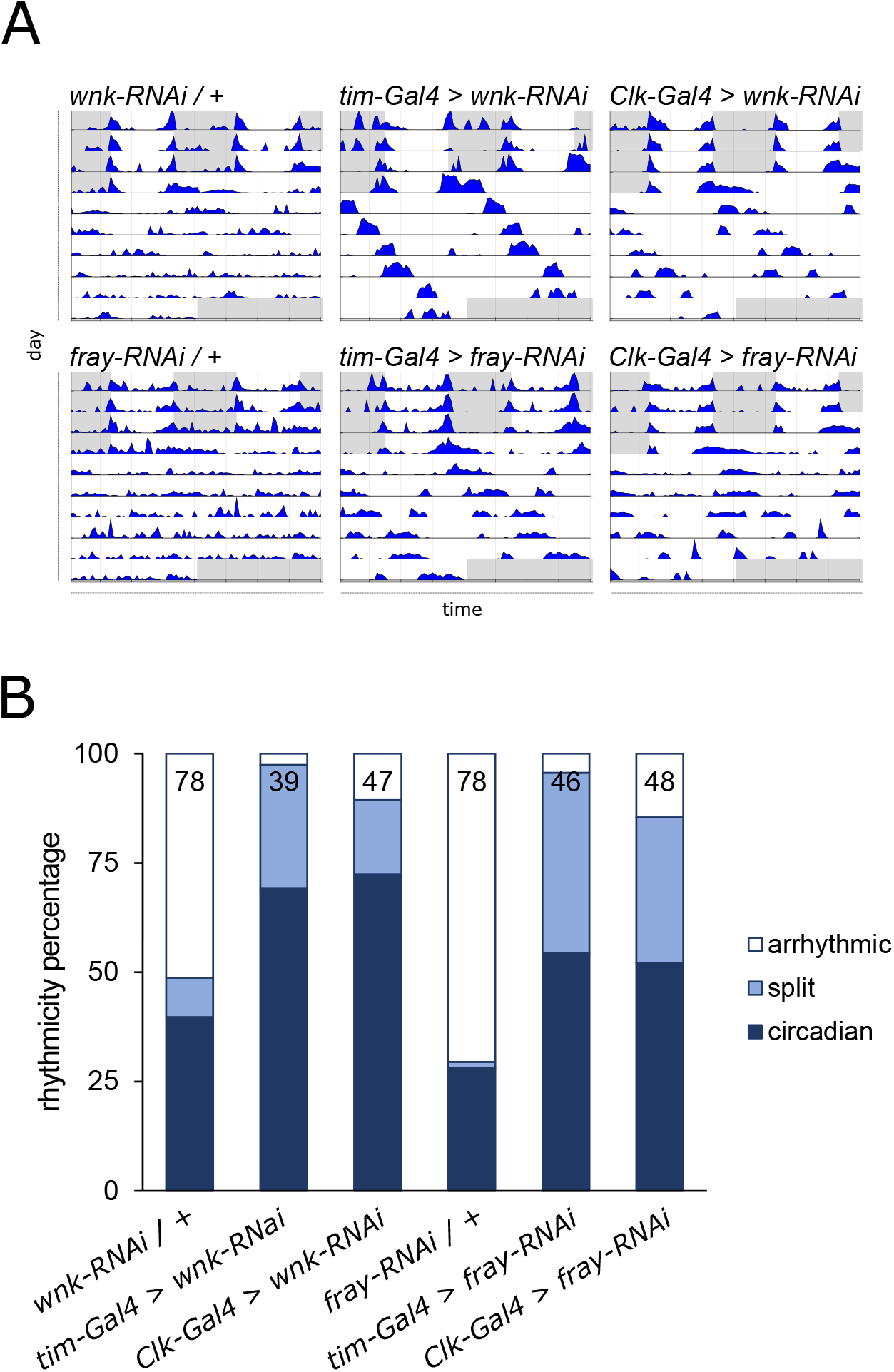
The kinases Wnk and Fray are required for normal light-input to the circadian clock. **(A)** Representative actograms of individual flies with altered *wnk* or *fray* expression and of individual control flies analyzed in LL as described in the legend for Figure 1. Flies express *wnk-RNAi* (*KK106928*) or *fray-RNAi* (*BL55878*) under the control of *tim-Gal4:27* or *Clk856-Gal4*. As controls, *Gal4* drivers (see Figure 1) and *UAS* lines were crossed to Canton S or *y w*. **(B)** Percentages of flies exhibiting circadian (dark blue), split (light blue) or arrhythmic (white) behavior in LL. Number of flies per genotype given inside bars.

### KCC and NKCC oppositely affect the GABA reversal potential

To determine if the behavioral effects of altered [Cl^−^]_i_ as a result of CCC modulation are reflected by physiological changes in the response to GABA, we performed patch-clamp recordings on the GABA responsive l-LNv neurons ^1,9–11^ to determine the GABA reversal potential (E_GABA_). The l-LNv express the ionotropic chloride channel receptor GABA_A_ ^9^, and first, we confirmed KCC and NKCC expression in these neurons using suitable intragenic *Gal4* insertions to drive a fluorescent cytoplasmic marker (Figure S1)^29^. In order to circumvent partial knock-down problems often associated with RNAi, we generated inducible knock-out lines under UAS control for *NKCC*, *kcc* and *qsm* using CRISPR technology ^12^ (see Methods). Similar to knock-down by RNAi, CRISPR-induced knock-out in all clock cells resulted in an increase of LL-rhythmicity for *NKCC* (26%), *kcc* (42%) and *qsm* (38%) (Table S1). Next, by measuring the GABA induced currents and calculating E_GABA_ from the resulting I–V curves, we confirmed that depletion of NKCC lowers E_GABA_ to more negative values than controls, while depletion of Qsm has the opposite effect (Figure 3, Table S2). Essentially the same results were previously obtained with *NKCC-RNAi* and *qsm-RNAi* flies ^1^, confirming the efficiency of the inducible CRISPR knock-out alleles and the consistency of the assay. As expected, *kcc^KO^*resulted in a more positive E_GABA_, which was higher than the resting membrane potential and thus essentially changing the polarity of GABA into an excitatory neurotransmitter (Figure 3B, green arrow in top trace). In contrast, overexpression of KCC drastically lowered E_GABA_, reaching even more negative values than that of *NKCC^KO^* flies (Figure 3D, Table S2).

**Figure 3:**
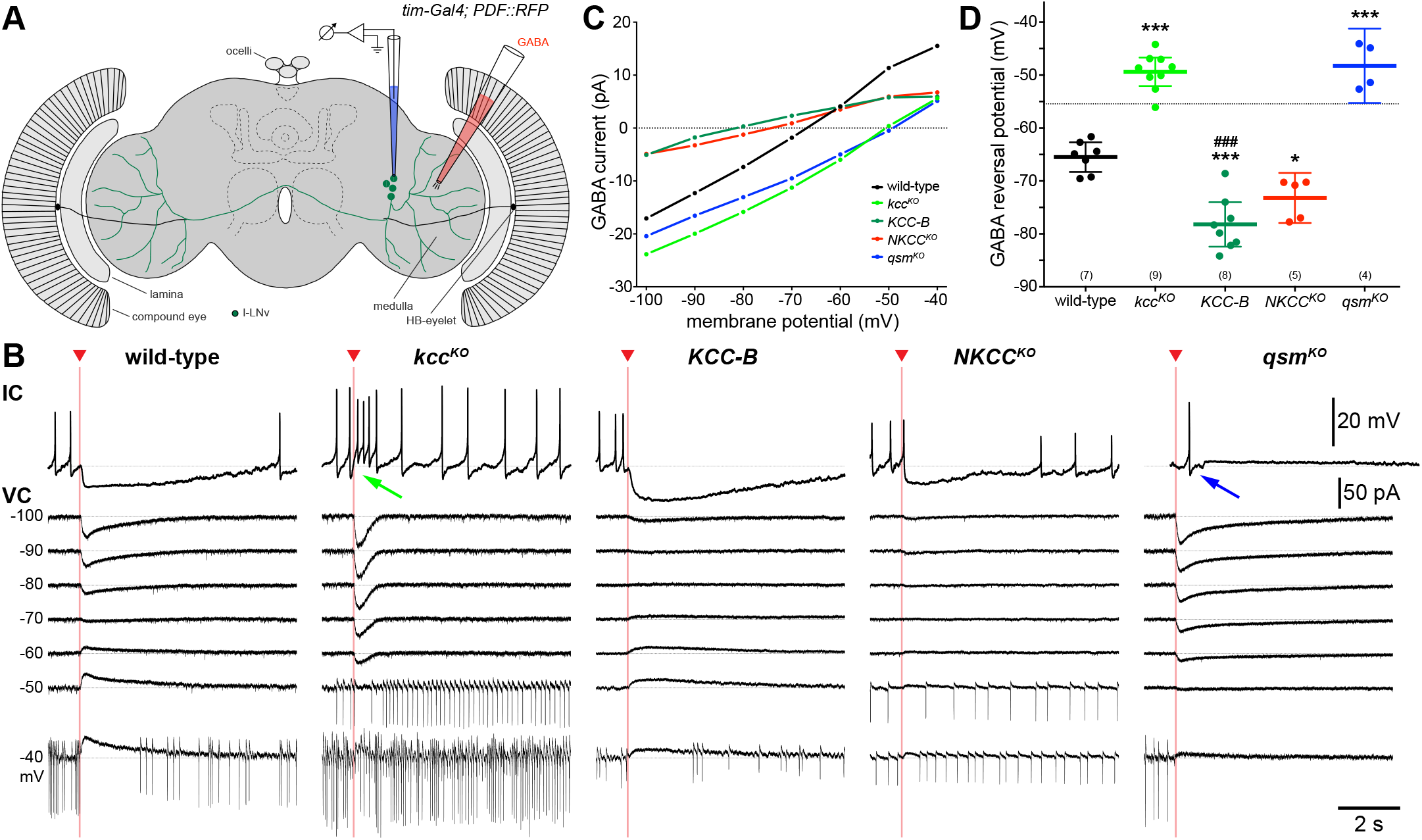
GABA responses of large LNv clock neurons with altered expression of Chloride Cotransporters. **(A)** Cartoon of the fly brain showing the recording setup with position and morphology of the l-LNv on one side, and other brain structures indicated for orientation. **(B)** Current clamp (IC, top at rest) and voltage clamp (VC, below at different holding potentials as indicated) whole-cell recordings of wild-type l-LNvs (*tim-Gal4:27; PDF::RFP*) and neurons with knock-out (*kcc^KO^*) or overexpression (*KCC-B*) of the ion transporter KCC as well as knock-out of *NKCC (NKCC^KO^*) and *qsm* (*qsm^KO^*) in response to brief GABA application (red arrowhead and line; pressure injection, 1-25 mM, 10-25 ms, 10 PSI). While control, *KCC-B* and *NKCC^KO^* neurons were hyperpolarized and inhibited, *kcc^KO^* and *qsm^KO^* neurons were depolarized and could even spike in response to GABA (see blue and green arrows). **(C)** Average currents evoked by GABA at the different holding potentials as measured from VC in (B) demonstrating a linear relationship with the currents reversing at different potentials for the indicated genotypes. **(D)** The GABA reversal potentials for *kcc^KO^* and *qsm^KO^* were above the average wild type resting membrane potential (−55.4 mV, indicated by dotted line) illustrating an excitatory effect for GABA in these neurons. Mean and confidence interval, N indicated in brackets; * p<0.05, *** and ### p<0.001; one-way ANOVA with Tukey’s *post hoc* test compared to wild-type controls (*) or between *kcc^KO^* and overexpression of *KCC-B* (#).

### KCC, NKCC and Qsm contribute to normal morning activity in long days

To test the relevance of CCC function and [Cl^−^]_i_ under more natural conditions, we tested flies with depleted KCC, NKCC, and Qsm levels under long days (16 h light : 8 h dark, 16:8 LD). Under these conditions, wild type *Drosophila melanogaster* shift their morning and evening activity peaks towards the lights-on and lights-off transition, and the l-LNv do play an important role in this adaptation ^13,14^. In normal 12 h : 12 h LD conditions *kcc^KO^, NKCC^KO^* and *qsm^KO^* flies exhibited wild type behavioral patterns comparable to the controls and were also robustly rhythmic in constant conditions (Figure S2, Table S3). In contrast, under 16:8 LD *kcc^KO^* showed a significant reduction of morning activity (ZT1 to ZT5) compared to the controls (Figure 4). Strikingly, *NKCC^KO^* had the opposite effect, with significant activity increase in the same time window (Figure 4). Surprisingly, *qsm^KO^* flies showed a similar activity increase as *NKCC^KO^*, although based on the LL behavioral results (Figure 1) and E_GABA_ measurements (Figure 3), a phenotype opposite to that of *NKCC^KO^* could have been expected. In any case, these experiments reveal a function for NKCC, KCC and Qsm in regulating behavioral activity in the morning of long days, with KCC promoting, and NKCC and Qsm repressing locomotor activity.

**Figure 4:**
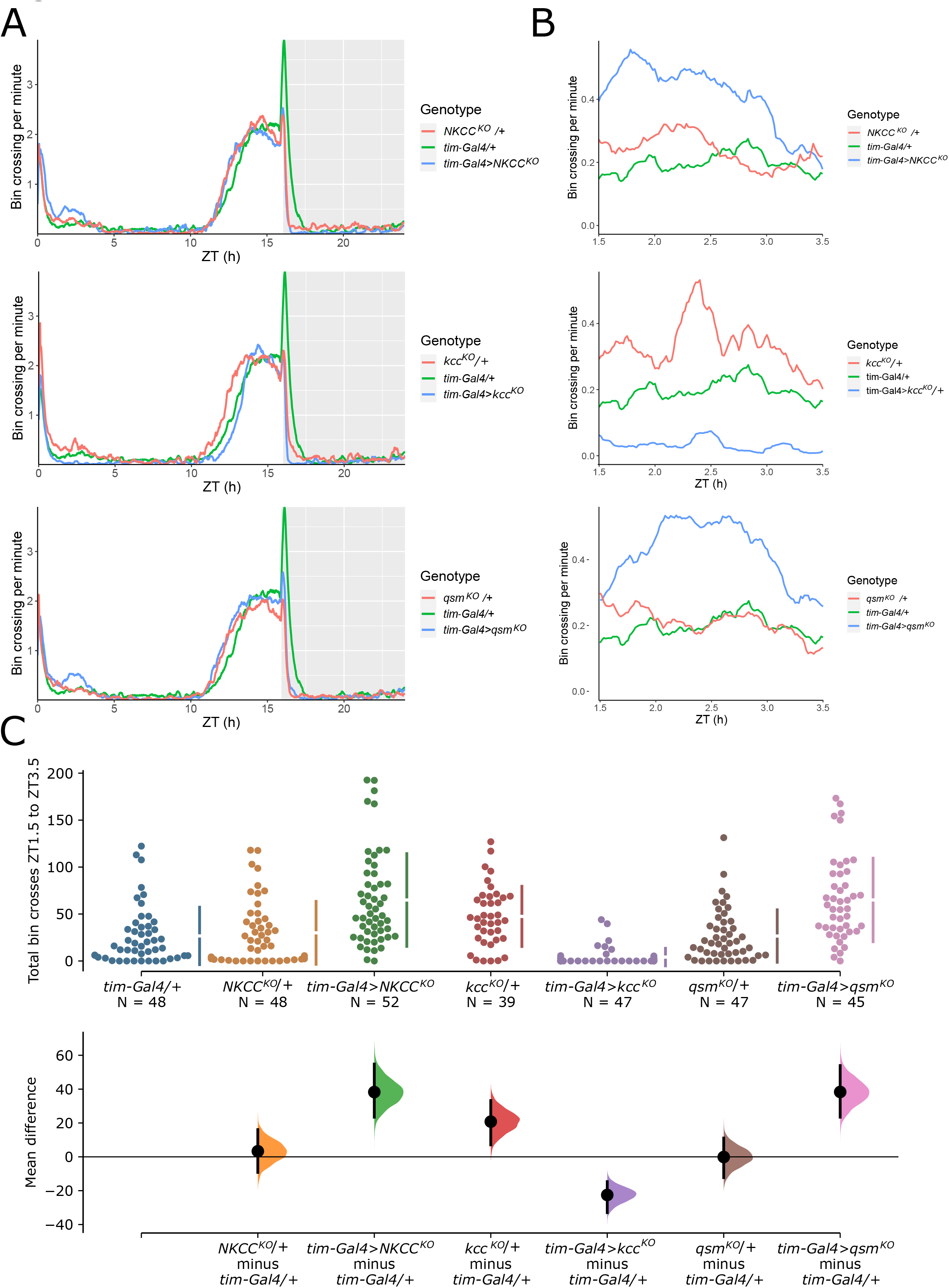
CCCs and *qsm* affect the morning peak in long days. **(A)** Smoothed average histograms of the last three days in 16 hr : 8 hr LD cycles for the indicated genotypes. **(B)** Activity of the different genotypes focusing on morning activity between ZT 1.5 and ZT 3.5. While knocking out *qsm* and *NKCC* leads to an activity increase, *kcc^KO^* flies reduce their activity compared to the controls. **(C)** Estimation statistics of the genotypes using *tim-Gal4/+* as reference (see Methods). The upper panel shows the total individual activity of each fly between ZT 1.5 and ZT 3.5 during the last three days of the 16 hr : 8 hr LD cycle, with the mean and standard error plotted as discontinuous line to the right. The lower panel shows the effect size (mean difference) of each of the genotypes compared to *tim-Gal4:27/+*. The controls *NKCC^KO^/+* and *qsm^KO^/+* do not show any difference to the control, while *tim-Gal4* driven knockout results in an activity increase (mean difference). In contrast, *kcc^KO^* in all clock cells leads to a reduction of activity, reversing the subtle activity increase of *kcc^KO^/+* control flies. The same comparison was performed to the *UAS* controls, showing similar results (not shown).

## Discussion

### CCCs regulate GABA polarity

CCCs are evolutionally conserved proteins that have been suggested to play a role within the circadian system of other insects and in mammals. In cockroaches, circadian neurons have been found to exert both inhibitory and excitatory responses to GABA, depending on the balance of NKCC1 / KCC2 activity. About 50% of all cockroach clock neurons respond to GABA, and about 30% of all GABA-responsive neurons show an increase of [Ca^2+^]_i_ as GABA response, which was correlated with enhanced NKCC expression compared to KCC ^15^. In the mammalian brain clock, the suprachiasmatic nucleus (SCN), expression and activity of the two opposing cotransporters NKCC1 and KCC2 underlie daily and regional variation. While NKCC1 is expressed both in the dorsal shell and the ventral core region, KCC2 can only be found in the latter ^16,17^. It has also been proposed that different environmental light conditions can affect the NKCC1 / KCC2 ratio, leading to temporal differences in [Cl^−^]_i_ and GABA polarity. However, the precise regulation of CCCs within the SCN and their contribution to modulating neuronal excitability in response to GABA remains elusive due to inconsistency between studies e.g., ^18–21^. Still, it is evident that GABA is widely expressed in the SCN ^22^ and appears to play an important role in synchronizing the circadian clock to light.

Here, we show that the response of the l-LNv clock neurons to GABA is also regulated by KCC and NKCC (Figure 3) ^1^, suggesting that GABA polarity within these neurons changes during the day. The observation that the l-LNv receive GABAergic input ^1,9–11^ further support this idea. Because both KCC and NKCC are expressed in subsets of other clock neuronal groups (LNd, DN1, DN3) (Figure S1) it is likely that GABA polarity changes throughout the clock network in a daily fashion. In addition to the l-LNv, expression of the GABA_A_ receptor has also been found in the accessory medulla, which contains dendritic processes from LNd and DN clock neurons, which like the l-LNv have been shown to depolarize in response to light ^9,23^.

### CCCs, Wnk, Fray and Quasimodo mediate light-input to circadian clock neurons

Alterations of KCC, NKCC, Wnk, Fray and Qsm expression within clock neurons results in rhythmic LL behavior (Figure 1, 2) ^1,2^, indicating a disruption of light-input to the clock. We previously demonstrated that NKCC and Qsm interact, whereby Qsm inhibits NKCC activity ^1^. Because Qsm expression is regulated by light and by the circadian clock ^2,24^, this suggests a light dependent-regulation of NKCC activity in clock neurons. In this model, light-activated Qsm would release NKCC inhibition, resulting in higher [Cl^−^]_i_ in the early morning or after light pulses potentially changing GABA polarity. These light-dependent changes of neuronal activity could contribute to both l-LNv mediated arousal responses (e.g. ^11,25^) as well as rhythmic LL-behavior mediated by LNd and DN clock neurons ^2,26–28^. This is further supported by *qsm*, *NKCC*, and *KCC* expression within subsets of l-LNv, LNd and DN (Figure S1) ^2,29^. The contribution of different clock neurons to LL-rhythmicity is also supported by the frequent observation of ‘split-rhythms’, which are characterized by the presence of two rhythmic components with different period lengths (Figure 1A, C) ^1^. Recent work suggests that the fly clock consists of independent units of neuronal oscillators, which in a decoupled state display different circadian periods ^30–32^. The combination of LL, which eliminates clock function in some of the units (e.g. the s-LNv), with manipulation of [Cl^−^]_i_ (which may not be equally effective in the different units depending on the *Gal4* driver and *RNAi* or overexpression line) could result in different coupling strength and oscillator function between and within the different clock neuronal groups, thereby explaining the emergence of split rhythms. Moreover, subtle differences between light intensities, for example due to different position within the incubator, could explain variation between individuals of the same genotype.

Interestingly, both constitutive reduction and increase of KCC and NKCC expression--and therefore presumably also of [Cl^−^]_i_--result in LL-rhythmicity (Figure 1, 2) ^1^. A possible explanation is the feedback regulation of NKCC and KCC activity by the Wnk/Fray kinases ^8^ (Figure 2). For example, overexpression of KCC is expected to lower [Cl^−^]_i_, which in turn activates the WNK/FRAY kinases leading to activation of NKCC and inhibition of KCC, thereby increasing [Cl^−^]_i_. Therefore, our findings indicate that any interference with the apparently intricate regulation of [Cl^−^]_i_ results in impaired neuronal activity and abnormal rhythmicity in LL, highlighting the importance for [Cl^−^]_i_ regulation within the neuronal circadian clock network for light-dependent behavior. Interestingly, [Cl^−^]_i_ regulation in a different subset of circadian clock neurons appears to be equally important for light-independent, purely circadian clock driven behavior (Schellinger et al 2021, this issue).

### Role of CCCs in seasonal adjustment to different day length

In mammals, switching of GABA polarity has been shown to not only contribute to phase delaying the circadian clock in response to a light pulse ^33^ but also to be implemented in adapting to seasonal changes in day length. For example, under long days, excitatory GABA responses were found to be increased in SCN neurons ^34,35^. The circadian network encodes photoperiod length by the phase relationship between core and shell clock neurons, which are in synchrony under short days and oscillate with an increased phase gap under long days. Transition between network states is suggested to be modulated by phase-dependent coupling of the two oscillators requiring GABA signaling ^35–37^. Interestingly, NKCC and KCC control of the GABA_A_ excitatory/inhibitory switch has been implicated in a range of neurodevelopmental and psychiatric disorders such as epilepsy, autism spectrum disorder and bipolar disorder, which have been reported to vary in a circadian and seasonal fashion ^38–41^.

Our results indicate that in fruit flies CCCs also contribute to the seasonable adjustment of day length. So far, the l-LNv have been implicated in the readjustment of the evening peak to long days ^13,14^. Here, we reveal a surprising influence of CCCs on the morning activity during long days, although the clock neuronal subset mediating this function remain elusive. NKCC and KCC depletion in all clock cells promoted or repressed activity, respectively (Figure 4). This effect is restricted to a 2-h time window immediately after the rapid decline of activity coinciding with lights-on, indicating that they are not simply light-driven and separate from the light-induced startle response. Interestingly, Qsm depletion also promoted activity, although we had expected an inhibitory effect. Nevertheless, the observation that Qsm function is also involved in regulating morning activity in long photoperiods, links this light-regulated protein again to NKCC and KCC function. Future work will show, which neurons contribute to this adjustment of morning activity during long photoperiods.

In summary, our work firmly links the regulation of [Cl^−^]_i_. and GABA polarity via the NKCC/KCC/Wnk/Fray feedback loop with light-input into the circadian clock and light-dependent behavior. Interference with [Cl^−^]_i_ results in abnormal rhythmic behavior in constant light and altered morning activity during long photoperiods and both behaviors also depend on the light-regulated membrane protein Qsm, which is known to regulate NKCC, and possibly also KCC activity. Regulation of [Cl^−^]_i_ and GABA polarity by CCCs therefore appears to be a conserved mechanism contributing to seasonal adjustment of the circadian clock and behavioral activity.

## Acknowledgements

We thank Dr. Aylin Rodan for sharing unpublished results and Mechthild Rosing for technical support. We thank members of our labs for comments on the manuscript. This work was supported a grant from the Deutsche Forschungsgemeinschaft given to RS (STA 421/8-1) and BBSRC grant (BB/W000865/1) to JJLH.

## Methods

### Flies

Flies were housed in plastic vials on standard fly food (0.7% agar, 1.0% soya flour, 8.0% polenta/maize, 1.8% yeast, 8.0% malt extract, 4.0% molasses, 0.8% propionic acid, 2.3% nipagen) under a 12 hr :12 hr LD cycle at 60% relative humidity. Fly stocks were kept at 18°C and crosses were reared at 25°C. Because the light sensitivity of wild type flies is influenced by the naturally occurring *s-tim/ls-tim* polymorphism ^42,43^, as well as by several mutations within the *jetlag* (*jet*) gene ^42,44^, all fly stocks used in this study were genotyped for *tim* and *jet* polymorphisms as described ^42,44^. If necessary *(Clk856-Gal4 and kcc-RNAi BL34584*), stocks were crossed into the *ls-tim* background. No stock carried one of the known mutant *jet* alleles (*jet^c^*, *jet^r^*,or *jet^set^*).

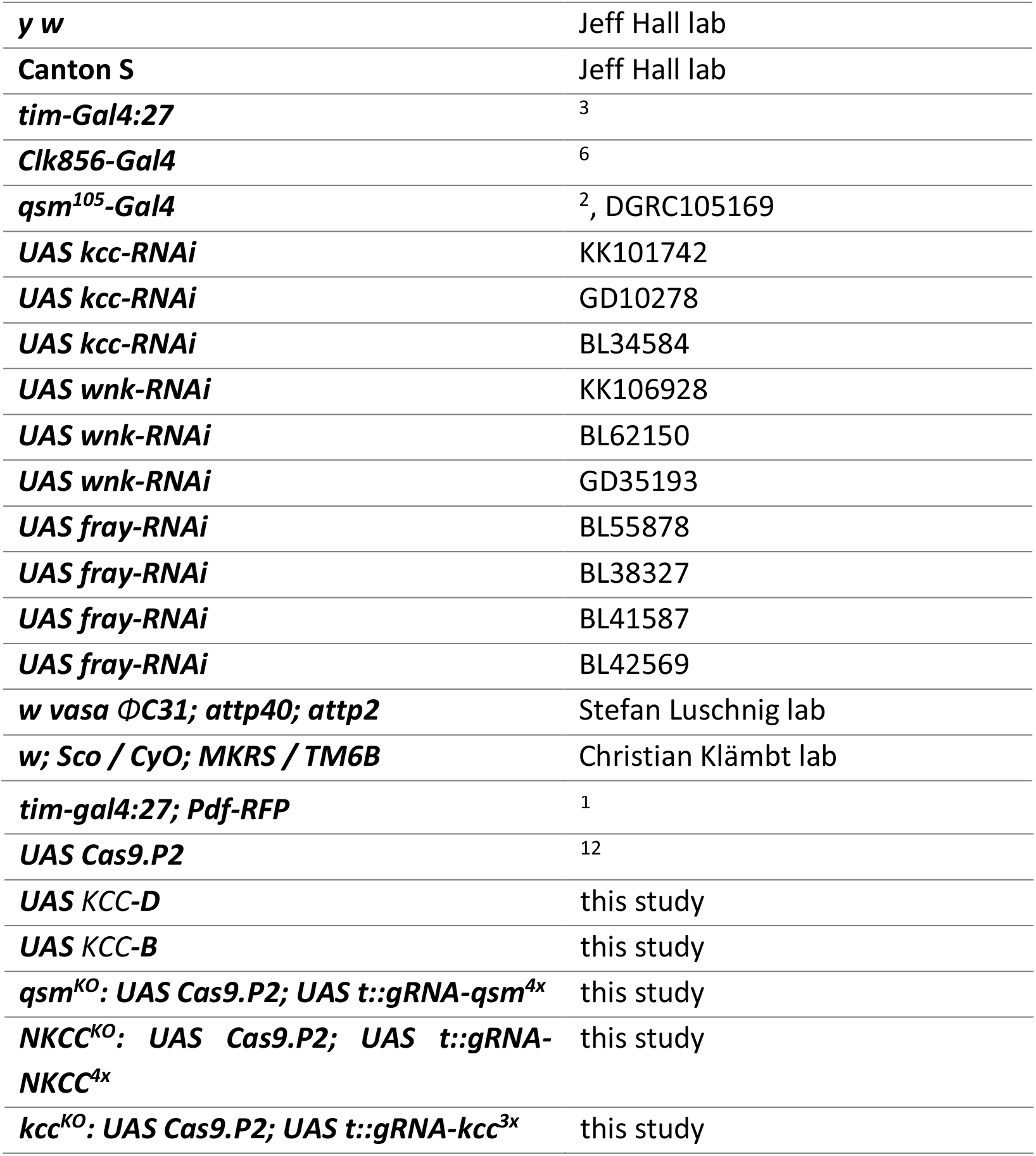

|  |  |
| --- | --- |
| **y w** | Jeff Hall lab |
| **Canton S** | Jeff Hall lab |
| ***tim-Gal4:27*** | 3 |
| ***Clk856-Gal4*** | 6 |
| ***qsm<sup>105</sup>-Gal4*** | 2, DGRC105169 |
| ***UAS kcc-RNAi*** | KK101742 |
| ***UAS kcc-RNAi*** | GD10278 |
| ***UAS kcc-RNAi*** | BL34584 |
| ***UAS wnk-RNAi*** | KK106928 |
| ***UAS wnk-RNAi*** | BL62150 |
| ***UAS wnk-RNAi*** | GD35193 |
| ***UAS fray-RNAi*** | BL55878 |
| ***UAS fray-RNAi*** | BL38327 |
| ***UAS fray-RNAi*** | BL41587 |
| ***UAS fray-RNAi*** | BL42569 |
| ***w vasa* ΦC31; *attp40*; *attp2*** | Stefan Luschig lab |
| ***w*; *Sco* / *CyO*; *MKRS* / *TM6B*** | Christian Klämbt lab |
| ***tim-gal4:27; Pdf-RFP*** | 1 |
| ***UAS Cas9.P2*** | 12 |
| ***UAS KCC-D*** | this study |
| ***UAS KCC-B*** | this study |
| ***qsm<sup>KO</sup>: UAS Cas9.P2; UAS t::gRNA-qsm<sup>4x</sup>*** | this study |
| ***NKCC<sup>KO</sup>: UAS Cas9.P2; UAS t::gRNA-NKCC<sup>4x</sup>*** | this study |
| ***kcc<sup>KO</sup>: UAS Cas9.P2; UAS t::gRNA-kcc<sup>3x</sup>*** | this study |

### Generation of UAS-KCC flies

Plasmids containing the full length *KCC-B* and *KCC-D* cDNAs were obtained from the Drosophila Genomic Resource Center (GH09271 for isoform B and LD02554 for isoform D). They were subcloned into pUAST-attB using BglII and XhoI for KCC-B (3.6kb) and NotI and XhoI for KCC-D (3.9kb), and transformed into competent Stellar Cells (Takara Bio). Recombinant plasmids were extracted using Roti-Prep Plasmid Mini (Carl Roth) and sent for sequencing using a general *pUAST* primer. The correct plasmids were then purified using the Plasmid Plus Purification Kit (Qiagen), injected into *vasaφ31;attP2;attP40*, and screened for red eyes.

### Generation of conditional CRISPR mutants

For conditional CRISPR knock-out of *qsm*, *NKCC* and *kcc*, fly lines were generated that express Cas9 as well as multiple gRNAs from a single tRNA::gRNA transcript under the control of the *Gal4/UAS* system ^12^. CRISPR Optimal Target Finder ^45^ was used to select four target gRNA sites each for *qsm* and *NKCC* and three target gRNA sites for *kcc*. Overlapping PCR primers encoding the gRNA sequences were ordered from Sigma-Aldrich and Eurofins Genomics. Primers were used on the pCFD6 template (Addgene #73915) using high-fidelity Phusion polymerase (NEB). Resulting PCR products were gel purified using Roti-Prep Gel Extraction (Carl Roth) and assembled with *Bbsl*-linearized pCFD6 backbone using NEBuilder HiFi DNA Assembly Master Mix (NEB) or In-Fusion HD Enzyme Premix (Takara Bio). The assembly mixture was transformed into competent DH5α or Stellar cells (Takara Bio) and transformants were screened in a colony PCR for inserts of the correct size using GoTaq polymerase (Promega) with pCFD6_F and pCFD6_R primers. Plasmids were extracted using Roti-Prep Plasmid Mini (Carl Roth) and send for sequencing with the pCFD6_F primer to GATC Eurofins Genomics. For transgenesis, plasmids were purified using Plasmid Plus Purification Kit (Qiagen) and injected into *w, vasa ΦC31; attp40; attP2* embryos. G0 offspring was batch crossed to *w; Sco/CyO; MKRS/TM6B* and F1 offspring was screened for red eyes and individually crossed to *w; Sco/CyO; MKRS/TM6B*. Flies carrying the construct on the third chromosome were combined with *UAS Cas9.P2* on the second chromosome.

Oligonucleotides:

QSM_PCR1_F
CGGCCCGGGTTCGATTCCCGGCCGATGCATGATCCTTTGACTCCGCATCGTTTCAGAGCTATGCTGG
AAAC
QSM_PCR1_R
GAGTAAGTACCTCGGCAGGCTGCACCAGCCGGGAATCGAACC
QSM_PCR2_F GCCTGCCGAGGTACTTACTCGTTTCAGAGCTATGCTGGAAAC
QSM_PCR2_R
TGCCACGGTCAACGTGTGTTTGCACCAGCCGGGAATCGAACC
QSM_PCR3_F
AACACACGTTGACCGTGGCAGTTTCAGAGCTATGCTGGAAAC
QSM_PCR3_R
ATTTTAACTTGCTATTTCTAGCTCTAAAACCGACTATTCCTGGCCAAGCGTGCACCAGCCGGGAATCGAACC
NKCC_PCR1_F
CGGCCCGGGTTCGATTCCCGGCCGATGCAGATTATAATTACGATCTCCGGTTTCAGAGCTATGCTGG
AAAC
NKCC_PCR1_R
GGGGTCCAATGGCAGCTCCGTGCACCAGCCGGGAATCGAACC
NKCC_PCR2_F
CGGAGCTGCCATTGGACCCCGTTTCAGAGCTATGCTGGAAAC
NKCC_PCR2_R
TGCTTAACGTGGCTGCAAAGTGCACCAGCCGGGAATCGAACC
NKCC_PCR3_F
CTTTGCAGCCACGTTAAGCAGTTTCAGAGCTATGCTGGAAAC
NKCC_PCR3_R
ATTTTAACTTGCTATTTCTAGCTCTAAAACGATCGCTCCTCTGATCTCCATGCACCAGCCGGGAATCGA
ACC
kcc_PCR1_F
TTCGATTCCCGGCCGATGCAGTCTCATAAGGTACCCTAATGTTTCAGAGCTATGCTGGAAAC
kcc_PCR1_R
TGCTAACCGCGATCTCGATGTGCACCAGCCGGGAATCGAACC
kcc_PCR2_F
CATCGAGATCGCGGTTAGCAGTTTCAGAGCTATGCTGGAAAC
kcc_PCR2_R
GCTATTTCTAGCTCTAAAACCATACACCGCTATTATCGAATGCACCAGCCGGGAATCGAACC
pCFD6_F
GTAGACATCAAGCATCGGTGG
pCFD6_R
TTAGAGCTTTAAATCTCTGTAGGTAG

### Behavior

Behavior experiments measuring locomotor activity were carried out using the Drosophila Activity Monitoring (DAM) system (TriKinetics). One to four days old male flies were loaded into small glass tubes, supplied with food (4% sucrose and 2% agar) and plugged with cotton. Monitors were kept in environmentally-controlled incubators (Percival Scientific) with white fluorescence light. All behavior experiments were performed at 25°C. For behavioral controls Gal4 drivers and UAS lines were crossed to Canton S or *y w*.

### LL and DD Behavior

To examine flies under constant conditions, the light regime was set to three or four days of 12 hr : 12 hr LD followed by seven days of either DD or LL. For LL experiments light intensity was set to ~100 lux. Data analysis was performed using the Fly-Toolbox in Matlab ^46^ that generates actograms and analyzes rhythmicity and period by autocorrelation, maximum entropy spectrum analysis (MESA) and periodogram. For behavior quantification of DD experiments, flies were manually scored as rhythmic or arrhythmic based on single actograms and rhythmicity analysis. Average period length was calculated from values given by autocorrelation analysis. For behavior quantification of LL experiments, flies were manually scored as rhythmic or arrhythmic based on single actograms and rhythmicity analysis. For this study, flies with an RS value ≥ 1.5 were considered rhythmic ^46^. Rhythmic LL-behavior was further classified as ‘circadian’ (if flies exhibited a single locomotor rhythm) or ‘split’ (if flies exhibited complex locomotor rhythms in which a long period was intersected by a shorter period).

### Long-day behavior

To study the behavior under long days, flies were subjected to four days of 12 hr: 12 hr LD cycles (100 lux) at 25°C. On the 5^th^ day, the light period was prolonged by four hours, two in the morning and two in the evening, resulting in a photoperiod of 16 hr light and 8 hr of darkness. Flies were in these conditions for seven days, and only the last three days were included in the analysis. Graphs were generated in R (R Foundation, Vienna, Austria), by plotting the average activity of each fly for each minute during the last three days. The experiments were repeated three times and the total average is shown in Figure 4. For the statistical analysis of the data, estimation statistics was used ^47^. Data were analyzed using DABEST ^48^, using the website available under https://www.estimationstats.com/#/. This approach offers a more informative way to analyze and interpret the results, since it focuses on the effect size and the precision of the experiment, and not only on significance testing. In order to compare the different groups of data at the same time, a shared-control plot was used, which is analogous to an ANOVA with multiple comparisons. The top part of the plot shows all the observed data points, with the mean and standard error for each of the genotypes to the right. The bottom part shows the effect size (mean difference) as a bootstrap 95% confidence interval (CI) with a bias corrected and accelerated (BCa) correction ^49^. The magnitude of the mean differences between groups can be easily compared to the reference, in this case *tim-Gal4/+*. The size of the CI shows the precision of the mean difference; the smaller it is, the more confident is the measure. The difference between datasets is visualized by the distance between the mean difference (black dot inside CI) and the reference line. If the CI is cutting the reference line, then both datasets may originate from the same distribution.

### Immunostaining

10 to 15 flies of each of the genotypes were fixed in 4% PFA for 2.5 hr at room temperature (RT). After fixation, the samples were washed six times with 0.1 M phosphate buffer (pH 7.4) with 0.1% Triton X-100 (PBS-T) at RT. The brains were dissected in PBS, then blocked with 5% normal goat serum in 0.5% PBS-T for 2 hr at RT and stained with pre-absorbed rabbit anti-PER (1:1000) ^50^ and mouse anti-GFP (Sigma G6519, 1:200) in 5% goat serum in 0.5% PBST for at least 48 hr at 4°C. After washing three times by PBS-T, the samples were incubated at 4°C overnight with goat anti-mouse AlexaFluor 488 nm (1:500) and anti-rabbit AlexaFluor 647 nm (Molecular Probes) in PBS-T. Brains were washed three times in PBS-T before being mounted in Vectashield. The images were taken using a Leica SP8 confocal microscope and processed using GIMP.

### Electrophysiology

Whole-cell current and voltage clamp recordings were performed on l-LNv visualized by RFP expression (*tim-Gal4; PDF::RFP*) and illumination with a 555 nm LED light source at ZT 1-3. Adult flies of either sex were collected two to five days post eclosion, briefly anaesthetized on CO_2_, decapitated and whole brains acutely dissected at room temperature (20-22°C) in extracellular saline containing (in mM): 101 NaCl, 1 CaCl_2_, 4 MgCl_2_, 3 KCl, 5 D-glucose, 1.25 NaH_2_PO_4_, 20.7 NaHCO_3_ and adjusted to a pH of 7.2. Cleaned brains were placed ventral side up in the recording chamber and neurons visualized using a 63x water-immersion lens on an upright microscope (Zeiss Examiner Z1). Glass electrodes with 8-15 MΩ resistance filled with intracellular solution (in mM: 102 K-gluconate, 17 NaCl, 0.94 EGTA, 8.5 HEPES, 0.085 CaCl_2_, 1.7 MgCl_2_ or 4 Mg·ATP and 0.5 Na·GTP, pH 7.2), an Axon MultiClamp 700B amplifier, an Axon DigiData 1440A digitizer (sampling rate: 20 kHz; filter: Bessel 10 kHz) and pClamp 10 and MultiClamp Commander software (Molecular Devices, Sunnyvale, CA, USA) were used for recordings. For voltage clamp recordings the capacitance and series resistance were corrected for and traces were offline low-pass filtered (8-pole Bessel, cut-off 1 kHz). GABA reversal potentials (E_GABA_) were calculated from the IV-curves resulting from the currents elicited by pressure injected GABA at different holding potentials in voltage clamp mode. For this, GABA (1-25 mM in extracellular saline) was injected (10-25 ms pulse) in the ipsilateral medulla via a glass pipette (3-6 MΩ) and a Picospritzer III (10 PSI; Parker Hannifin, NH, USA). The liquid junction potential was calculated as 13 mV and subtracted from all the membrane voltages. Values are given as mean and 95% confidence interval. The Kolmogorov-Smirnov test was used to test for normality and a 1-way ANOVA followed by Tukey’s multiple comparison test for differences between wild-type and experimental neurons. The statistical tests were performed using Prism (GraphPad Software Inc., La Jolla, CA, USA) and the figure arranged in Illustrator (Adobe Systems Inc., San Jose, CA, USA).

---

**Figure S1:**
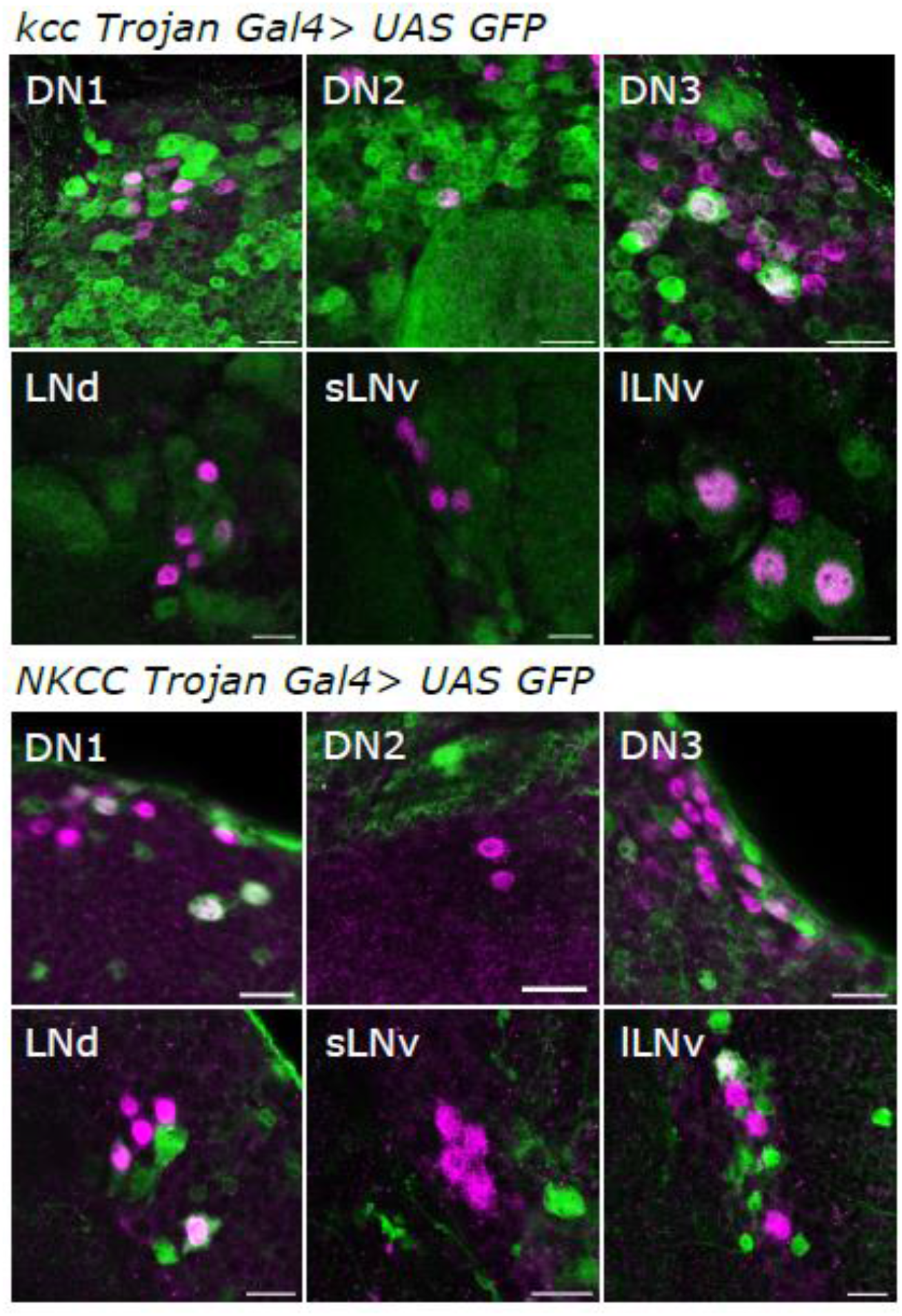
KCC and NKCC expression in the clock cells. UAS GFP S65T was expressed using the available Trojan Gal4 lines (BL66858 for *kcc* and BL77185 for *NKCC*) and visualized with a GFP antibody in green and counterstained with PER in magenta. *kcc-Gal4* is expressed in the 4 l-LNv, 3 LNd, 7-9 DN3, 2 DN2, 4 DN1p and 2 DN1a. *NKCC-Gal4* is expressed in 1 l-LNv, 3 LNd, 5 DN3, 5 DN1p, 2 DN1a. To test if the 5^th^ s-LNv was stained, we counterstained the brains with PDF, and the only colocalization observed was in the l-LNv for both Trojan Gal4 lines (not shown). For each genotype a total of 6 brains were analyzed. Scale bar=10μm

**Figure S2:**
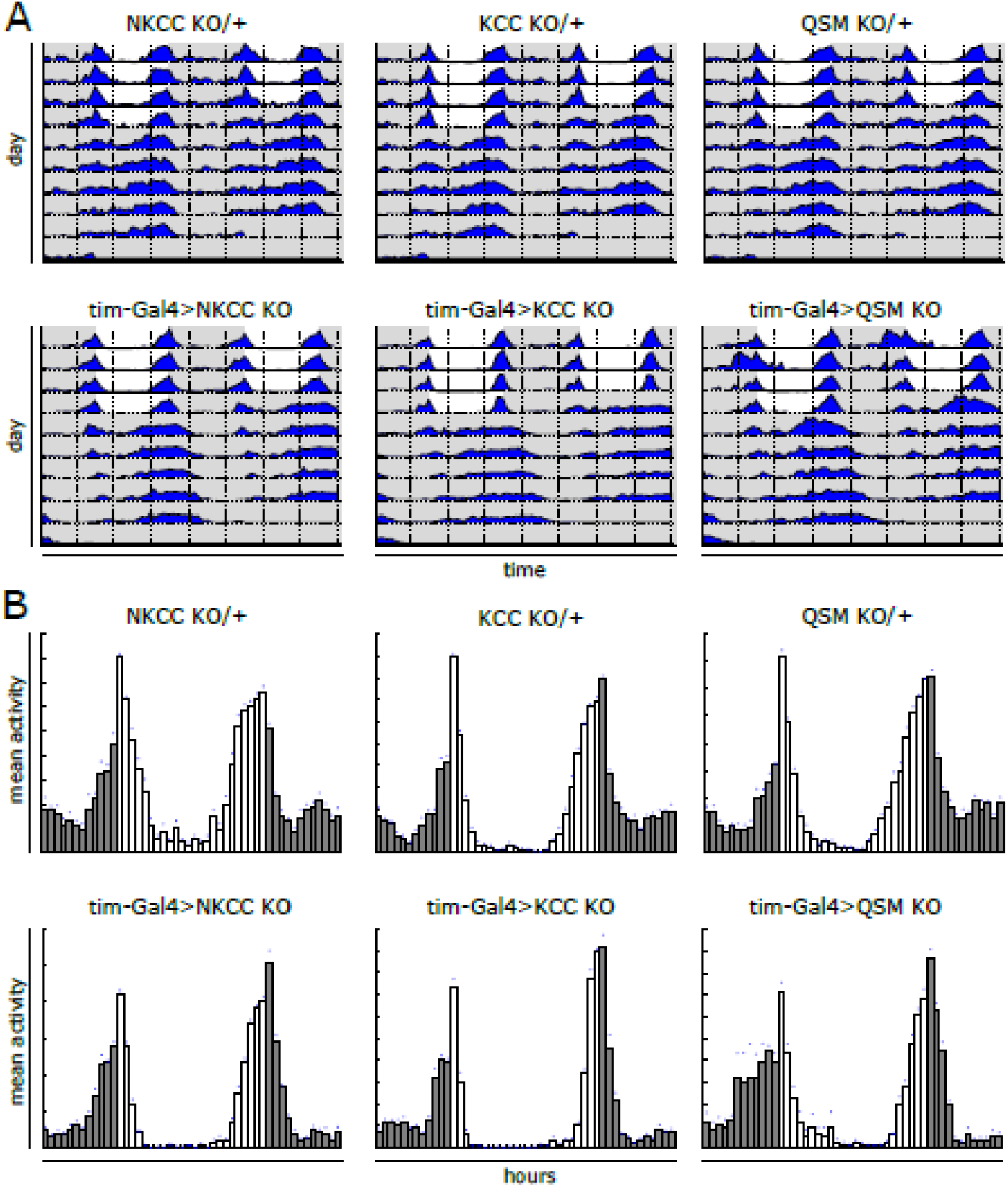
Normal behavior of knock-out lines in LD and DD. Representative average actograms **(A)** for the entire experiment (LD and LL) and histograms **(B)** for the LD part (n=16) of the genotypes indicated. Flies were exposed to four days of 12:12hr LD, followed by five days of constant darkness at 25°C. White areas and bars indicate lights-on, grey areas and bars lights-off.

**Table S1:** Quantification of behavioral rhythmicity in constant light (LL).

| **Genotype** | **% rhythmic** | **% circadian** | **% split** | **% arrhythmic** | **n** |
| --- | --- | --- | --- | --- | --- |
| *tim-Gal4:27 / +* | 57.0 | 45.0 | 12.0 | 43.0 | 100 |
| *Clk856-Gal4 / +* | 37.6 | 28.0 | 9.7 | 62.4 | 93 |
| *qsm<sup>105</sup>-Gal4 / +* | 44.3 | 41.0 | 3.3 | 55.7 | 61 |
| *kcc-RNAi KK101742 / +* | 57.4 | 39.4 | 18.1 | 42.6 | 54 |
| *tim-Gal4:27 > kcc-RNAi KK101742* | 100.0 | 68.8 | 31.3 | 0.0 | 48 |
| *Clk856-Gal4 > kcc-RNAi KK101742* | 89.1 | 47.8 | 41.3 | 10.9 | 41 |
| *qsm<sup>105</sup>-Gal4 > kcc-RNAi KK101742* | 80.9 | 57.4 | 23.4 | 19.1 | 38 |
| *kcc-RNAi GD10278 / +* | 37.6 | 33.3 | 4.3 | 62.4 | 93 |
| *tim-Gal4:27 > kcc-RNAi GD10278* | 86.7 | 62.2 | 24.4 | 13.3 | 45 |
| *Clk856-Gal4 > kcc-RNAi GD10278* | 70.2 | 38.3 | 31.9 | 29.8 | 47 |
| *qsm<sup>105</sup>-Gal4 > kcc-RNAi GD10278* | 79.5 | 68.2 | 11.4 | 20.5 | 44 |
| *kcc-RNAi BL34584 / +* | 37.7 | 33.8 | 3.9 | 62.3 | 77 |
| *tim-Gal4:27 > kcc-RNAi BL34584* | 90.3 | 48.4 | 41.9 | 9.7 | 31 |
| *Clk856-Gal4 > kcc-RNAi BL34584* | 46.8 | 44.7 | 2.1 | 53.2 | 47 |
| *qsm<sup>105</sup>-Gal4 > kcc-RNAi BL34584* | 56.3 | 50.0 | 6.2 | 43.8 | 16 |
| *KCC-D / +* | 7.4 | 7.4 | 0.0 | 92.6 | 27 |
| *tim-Gal4:27 > KCC-D* | 96.3 | 81.5 | 14.8 | 3.7 | 27 |
| *Clk856-Gal4 > KCC-D* | 73.9 | 52.2 | 21.7 | 26.1 | 23 |
| *qsm<sup>105</sup>-Gal4 > KCC-D* | 50.0 | 43.8 | 6.3 | 50.0 | 16 |
| *KCC-B / +* | 2.7 | 2.7 | 0.0 | 97.3 | 37 |
| *Clk856-Gal4 > KCC-B* | 59.5 | 48.6 | 10.8 | 40.5 | 37 |
| *qsm<sup>105</sup>-Gal4 > KCC-B* | 36.4 | 27.3 | 9.1 | 63.6 | 11 |
| *wnk-RNAi KK106928 / +* | 48.7 | 39.7 | 9.0 | 51.3 | 78 |
| *tim-Gal4:27 > wnk-RNAi KK106928* | 97.4 | 69.2 | 28.2 | 2.6 | 39 |
| *Clk856-Gal4 > wnk-RNAi KK106928* | 89.4 | 72.3 | 17.0 | 10.6 | 47 |
| *wnk-RNAi GD35193 / +* | 38.3 | 31.9 | 6.4 | 61.7 | 47 |
| *tim-Gal4:27 > wnk-RNAi GD35193* | 86.7 | 60.0 | 26.7 | 13.3 | 15 |
| *Clk856-Gal4 > wnk-RNAi GD35193* | 91.5 | 48.9 | 42.6 | 8.5 | 47 |
| *fray-RNAi BL55878 / +* | 29.5 | 28.2 | 1.3 | 70.5 | 78 |
| *tim-Gal4:27 > fray-RNAi BL55878* | 95.7 | 54.3 | 41.3 | 4.3 | 46 |
| *Clk856-Gal4 > fray-RNAi BL55878* | 85.4 | 52.1 | 33.3 | 14.6 | 48 |
| *fray-RNAi BL38327 / +* | 34.2 | 28.9 | 5.3 | 65.8 | 76 |
| *tim-Gal4:27 > fray-RNAi BL38327* | 100.0 | 55.3 | 44.7 | 0.0 | 47 |
| *Clk856-Gal4 > fray-RNAi BL38327* | 66.0 | 46.8 | 19.1 | 34.0 | 47 |
| *fray-RNAi BL41587 / +* | 51.9 | 44.2 | 7.8 | 48.1 | 77 |
| *tim-Gal4:27 > fray-RNAi BL41587* | 93.6 | 55.3 | 38.3 | 6.4 | 47 |
| *Clk856-Gal4 > fray-RNAi BL41587* | 62.5 | 45.8 | 16.7 | 37.5 | 48 |
| *fray-RNAi BL42569 / +* | 35.9 | 30.8 | 5.1 | 64.1 | 78 |
| *tim-Gal4:27 > fray-RNAi BL42569* | 75.0 | 54.5 | 20.5 | 25.0 | 44 |
| *Clk856-Gal4 > fray-RNAi BL42569* | 65.0 | 40.0 | 25.0 | 35.0 | 40 |
| *qsm<sup>KO</sup> / +* | 62.5 | 56.3 | 6.3 | 37.5 | 16 |
| *tim-Gal4:27 > qsm<sup>KO</sup>* | 100.0 | 75.0 | 25.0 | 0.0 | 16 |
| *NKCC<sup>KO</sup> / +* | 66.7 | 60.0 | 6.7 | 33.3 | 15 |
| *tim-Gal4:27 > NKCC<sup>KO</sup>* | 92.3 | 53.8 | 38.5 | 7.7 | 13 |
| *kcc<sup>KO</sup> / +* | 58.1 | 45.2 | 12.9 | 41.9 | 31 |
| *tim-Gal4:27 > kcc<sup>KO</sup>* | 100.0 | 55.2 | 44.8 | 0.0 | 29 |

**Table S2:** GABA reversal potential in control and experimental large LNv.

| Genotype | E<sub>GABA</sub> (mV) | N | p-value (*) | p-value (#) |
| --- | --- | --- | --- | --- |
| *PDF::RFP; tim-gal4:27 / +* | **-65.5** (-68.3, -62.7) | 7 |  |  |
| *PDF::RFP; tim-gal4:27 > kcc<sup>KO</sup>* | **-49.4** (-52.1, -46.7) | 8 | < 0.001 |  |
| *PDF::RFP; tim-gal4:27 > KCC-B* | **-78.2** (-82.4, -74.0) | 9 | < 0.001 | < 0.001 |
| *PDF::RFP; tim-gal4:27 > NKCC<sup>KO</sup>* | **-73.2** (-77.9, -68.5) | 5 | 0.02 |  |
| *PDF::RFP; tim-gal4:27 > qsm<sup>KO</sup>* | **-48.3** (-55.3, -41.2) | 4 | < 0.001 |  |

For each genotype the mean and 95% CI (lower, higher) for the GABA reversal potential (E<sub>GABA</sub>), the number of recorded neurons (N) and the p-value of the one-way ANOVA with Tukey's *post hoc* test compared to wild-type *PDF::RFP; tim-gal4:27 / +* control (\*) or between *PDF::RFP; tim-gal4:27 > kcc<sup>KO</sup>* and *PDF::RFP; tim-gal4:27 > KCC-B* (#) are given.

**Table S3.**
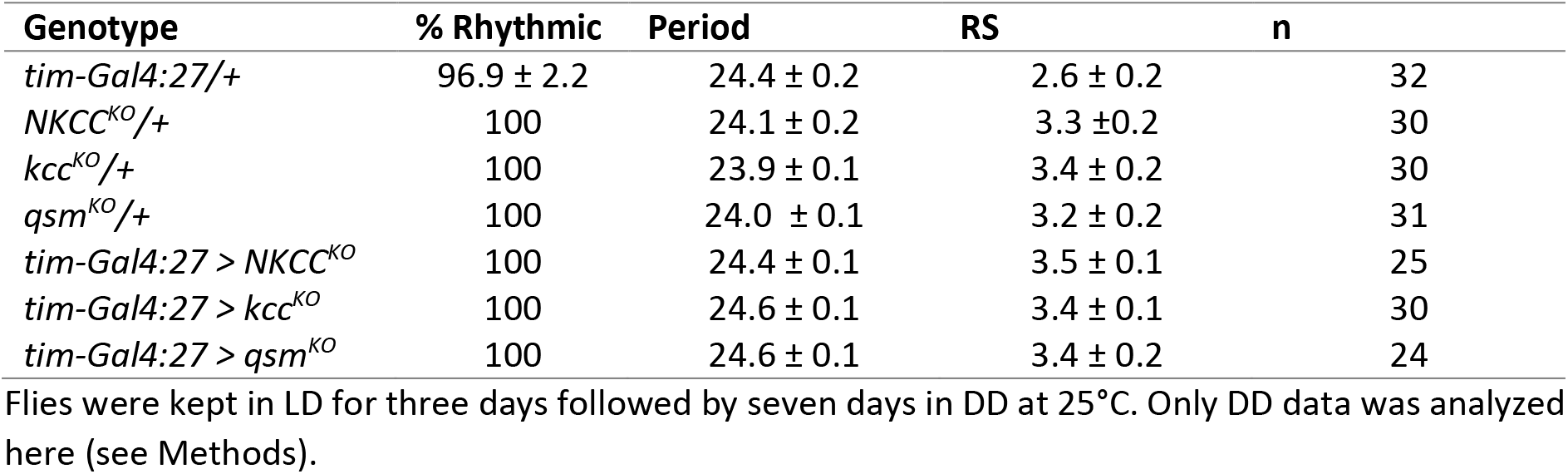
Free running behaviour of the knock-outs. All values represent the average ± SEM.

